# Automated analysis of rabbit knee calcified cartilage morphology using micro-computed tomography and deep learning segmentation

**DOI:** 10.1101/2020.08.21.260992

**Authors:** Santeri J. O. Rytky, Lingwei Huang, Petri Tanska, Aleksei Tiulpin, Egor Panfilov, Walter Herzog, Rami K. Korhonen, Simo Saarakkala, Mikko A. J. Finnilä

**Affiliations:** Research Unit of Medical Imaging, Physics and Technology, University of Oulu, Oulu, Finland; Department of Applied Physics, University of Eastern Finland, Kuopio, Finland; Department of Diagnostic Radiology, Oulu University Hospital, Oulu, Finland; Ailean Technologies Oy, Oulu, Finland; Human performance laboratory, Faculty of Kinesiology, University of Calgary, AB, Calgary, Canada

**Keywords:** Bone μCT, Bone histomorphometry, Animal models, Osteoarthritis

## Abstract

**Purpose:** Only little is known how calcified cartilage (CC) structure changes during exercise, aging and disease. CC thickness (CC.Th) can be analyzed using conventional histological sections. Micro-computed tomography (μCT) allows for three-dimensional (3D) imaging of mineralized tissues, however, the segmentation between bone and CC is challenging. Here, we present state-of-the-art deep learning segmentation for μCT images to enable assessment of CC morphology.

**Methods:** Sixteen knees from twelve New Zealand White rabbits were dissected into osteochondral samples from six anatomical regions: lateral and medial femoral condyles, lateral and medial tibial plateaus, femoral groove and patella (*n* = 96). Samples were imaged with μCT and processed for conventional histology. Manually segmented CC from the histology and reconstructed μCT images was used as the gold standard to train segmentation models with different encoder-decoder architectures. The models with the greatest out-of-fold evaluation Dice score were used for automated CC.Th analysis. Subsequently, the automated CC.Th analysis was compared across a total of 24 regions, co-registered between the imaging modalities, using Pearson correlation and Bland-Altman analyses. Finally, the anatomical variation in CC.Th was assessed via a Linear Mixed Model analysis.

**Results:** The best segmentation models yielded average Dice scores of 0.891 and 0.807 for histology and μCT segmentation, respectively. The correlation between the co-registered regions across the modalities was strong (*r* = 0.897). The Bland-Altman analysis yielded a bias of 21.9 μm and a standard deviation of 21.5 μm between the methods. Finally, both methods could separate the CC morphology between the patella, femoral, and tibial regions (*p* < 0.001).

**Conclusion:** The presented method allows for *ex vivo* 3D assessment of CC.Th in an automated and non-destructive manner. We demonstrated its utility by quantifying CC.Th in different anatomical regions. CC.Th was the thickest in the patella and the thinnest in the tibial plateau.

**Graphical abstract:** 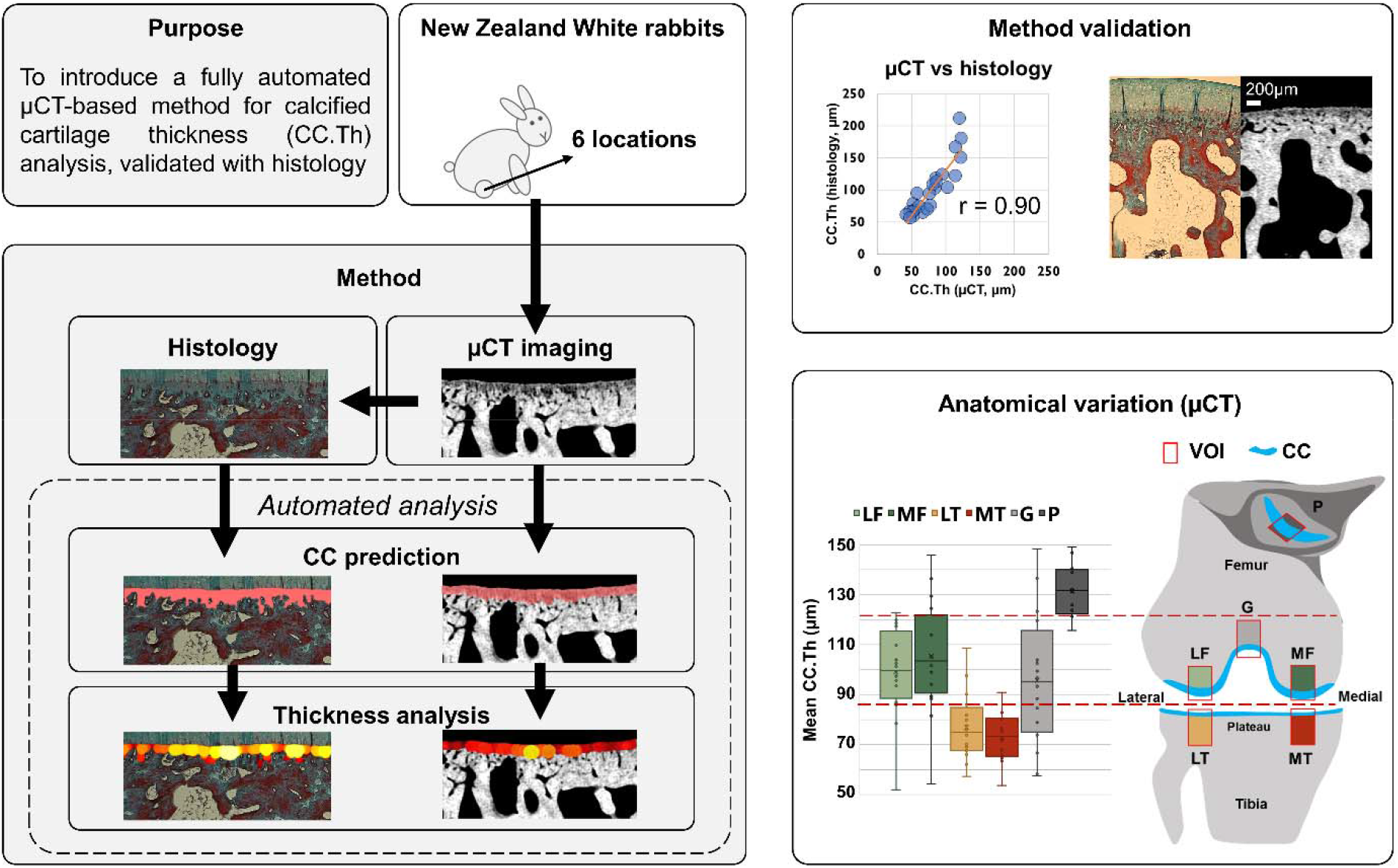

We present a μCT-based method with deep learning segmentation for analyzing calcified cartilage thickness (CC.Th). The method is compared throughout the study against conventional histology. The comparison against co-registered regions yielded a strong Pearson correlation (r = 0.90). Both methods were able to separate the CC.Th properties between tibia, femur, and patella.

## Introduction

Calcified cartilage (CC) is a mineralized tissue delineated from the non-calcified articular cartilage by the tidemark, and from the subchondral bone by the cement line^(1)^. The CC has an important role in anchoring the articular cartilage to the subchondral bone via individual collagen fibrils^(2)^. For healthy conditions, the relative CC thickness (CC.Th) to the total cartilage is nearly constant, but the CC volume relative to the total cartilage volume varies and has been shown to range from 3.23% to 8.8%^(3)^. Blood vessels from the subchondral bone extend into the CC layer, providing nutrients to the local chondrocytes^(1)^. Furthermore, based on the current literature, CC is a dynamic tissue undergoing changes with mechanical loading, aging and joint pathology, *e.g.* osteoarthritis^(4)^.

The thickness of articular cartilage^(5–6)^ and subchondral bone^(7)^ varies greatly in different areas of the knee joint with a high thickness in heavily loaded areas. It can be hypothesized that similar changes are present in the CC as well. Early study on CC.Th revealed regional differences within the human femoral head^(3)^. Furthermore, clear regional differences in equine CC have been reported^(8–9)^. In contrast, in canine knees only minor regional differences have been found^(5)^. These differences related to anatomical location could be linked to the local loading environment.

In general, exercise and loading are thought to affect the CC structure. The intensity of exercise on heavily loaded joint regions is associated with thicker CC in equine tarsi^(10)^ and carpus, even without changes in the overlying non-calcified cartilage^(11)^. An increase in the canine CC.Th was observed with high-intensity exercise^(12)^. In contrast, unloading of knees with immobilization resulted in thinner CC in canine knees^(13)^. In the human knee joint, similar findings have been reported; both articular and calcified cartilage are thick in load-bearing areas and thin under the menisci of the knee^(14)^.

Two competing events occur in aging CC: calcification of the deep articular cartilage via advancement of the tidemark^(15)^ and endochondral ossification (bone replacing CC at the cement line)^(16)^. The latter is likely dominant since aging accelerates the thinning of CC and increases the number of tidemarks^(16–17)^. Although CC.Th varies across humans and different animal species^(18)^, similar changes in aging CC have been found in animal models. Thinning of CC, increases in vessel invasion^(19)^, as well as chondrocyte apoptosis^(20)^ have been reported in murine CC with aging. On the other hand, *Murray et al.* reported an age-related increase in CC.Th in the equine tarsometatarsal joint^(21)^. Joint pathology can also induce tissue responses in the CC. Remodeling of CC^(16–17)^ occurs during OA progression, contributing to a decrease in articular cartilage thickness^(22)^. Microfractures in the CC, subchondral bone plate, and the trabeculae, lead to the formation of cysts and channels, thereby affecting the cross-talk between articular cartilage and subchondral bone^(1)^.

Traditionally, CC imaging has been performed on images obtained from histological sections^(3)^ as well as backscattered scanning electron microscopy (SEM) in equine^(16)^ and human joints^(23–24)^. Both histology and SEM require extensive and time-consuming sample processing protocols, and allow for two-dimensional (2D) imaging only. Nowadays, three-dimensional (3D) volumetric reconstruction of histological^(25)^ and SEM images^(26)^ is possible with serial sectioning and imaging, but the associated processing is laborious and has the potential to introduce errors.

Micro-computed tomography (μCT) has been widely used to characterize 3D morphology in micron level, including CC^(27–28)^. In contrast to histology and SEM, only minimal sample processing is required in μCT. We showed previously that μCT images of the human subchondral plate contain both the mineralized CC and the subchondral bone^(29)^. Indeed, CC cannot be separated from bone with low-resolution μCT imaging but becomes visible only in high-resolution μCT images^(30)^. However, because of the very minor difference in mineralization between the subchondral bone and CC, it is challenging to delineate the interface between CC and subchondral bone also in high-resolution μCT imaging.

The identification of the tidemark and cement line from μCT images is often conducted manually by researchers. This is a subjective and highly time-consuming endeavor, especially for tissues with complex shapes. Deep convolutional neural networks (CNNs) have recently shown great promise for automating various segmentation problems. U-Net^(31)^ has been the most popular segmentation architecture for biomedical images in recent years, and it has also been applied to μCT data^(32)^. However, the newly introduced Feature Pyramid Networks (FPN) allow for capturing both low-resolution global features as well as high-resolution local features at a low computational cost^(33)^. Conventional training of CNNs is conducted by initializing the coefficients from a random distribution. An alternative training approach is transfer learning, in which the network is initialized from an existing model, often pre-trained on ImageNet dataset^(34–35)^. Notably, such approach works efficiently across domains beyond natural images^(36–37)^. For example, transfer learning from deep residual networks^(38)^ has been used to classify pulmonary nodules from CT images^(39)^, or segment the lungs in chest X-rays^(40)^.

In this study, we propose an accurate framework for automated μCT-based evaluation of the CC.Th in 3D. This requires introducing state-of-the-art deep learning architectures for CC segmentation. To demonstrate the validity of the method, we perform direct comparison of CC.Th between μCT and conventional histology. We utilized osteochondral samples of New Zealand White rabbits, a frequently used animal model for various musculoskeletal diseases. Furthermore, we hypothesize that the CC.Th varies in different anatomical locations of the knee. We demonstrate the capability of our automatic framework by assessing differences in CC.Th between the different anatomical locations.

## Materials and Methods

### Sample collection

Sixteen knees were collected from twelve healthy, skeletally mature female New Zealand White rabbits (strain 052 CR). Eight knees were collected from four rabbits (age: 14 months) and eight knees from eight rabbits (age: 12.5 months). Each knee was dissected and divided into six anatomical regions: lateral and medial femoral condyle, lateral and medial tibial plateau, femoral groove and patella (n = 96, Table 1). Details on animal housing, husbandry conditions and diet are detailed in a previous study^(41)^. All experiments were carried out under the guidelines of the Canadian Council on Animal Care and were approved by the committee on Animal Ethics at the University of Calgary (Renewal 3 for ACC Study #AC11-0035).

**Table 1.** Descriptive statistics of the rabbits used in the study. On the right, the number of images and samples (separated by / mark) segmented manually is described. These segmentations are used as training data for the deep learning models.

| # animals | # knees | # Samples | # Histology slices | Manual segmentations |  |
| --- | --- | --- | --- | --- | --- |
|  |  |  |  | Histology | μCT |
| 12 | 16 | 96 | 3 / sample | 253 / 87 | 1050 / 60 |

### Imaging

The dissected osteochondral samples were formalin-fixed. Prior to imaging, samples were wrapped in moist paper, and placed in plastic vials (Cryo.s™) for positional stability. The samples were subsequently imaged using a desktop μCT scanner (Skyscan 1272, Bruker microCT, Kontich, Belgium) with a tube voltage of 50 kV, current of 200 μA, and a 0.5 mm aluminum filter. The scanning was conducted in a step of 0.2° over 360° and finally, 1800 projection images with an isotropic pixel size of 3.2 μm were obtained.

The images were reconstructed using the manufacturer’s software (NRecon, version 1.7.0.4). A narrow window with attenuation coefficients 0.085-0.141 was used to provide high contrast between the bone and CC. The volumes-of-interest (VOI) of all samples were selected from the central load-bearing area (VOI size = 2 mm × 2 mm × sample height). This selection reduced the μCT image stacks to a reasonable size (from ~12 GB to ~700 MB per sample) for the subsequent analysis. See Supplementary Figure 1 for examples of the preprocessing steps.

After the μCT imaging, samples were prepared for histological analysis. Samples were decalcified using a standard protocol (with EDTA solution), paraffin-embedded, and cut into 5-μm-thick sections using a microtome (three sections from each region). The sections were stained with Masson-Goldner’s trichrome for identification of the CC layer and imaged with a light microscope (Axioimager 2; Carl Zeiss MicroImaging Gmbh, Jena, Germany; control software = AxioVision; resolution = 2.56 μm). A total of 281 sections were used in this study.

### Training CC segmentation models

For the histology images, 253 continuous CC structures were segmented manually from 87 samples based on the distinct collagen staining in CC against articular cartilage and subchondral bone (Figure 1a). For the μCT, manual annotation was conducted for 60 samples from 10 knees according to two criteria: 1) CC region with a distinct grayscale gradient and 2) presence of chondrocytes inside the CC layer (Figure 1c, blue arrows). Annotations were done for 10-30 slices per sample, evenly spaced within each volume. The manual annotations were used as the gold standard for the automated segmentation algorithms and for conducting a reference analysis for the CC morphology.

**Figure 1.**
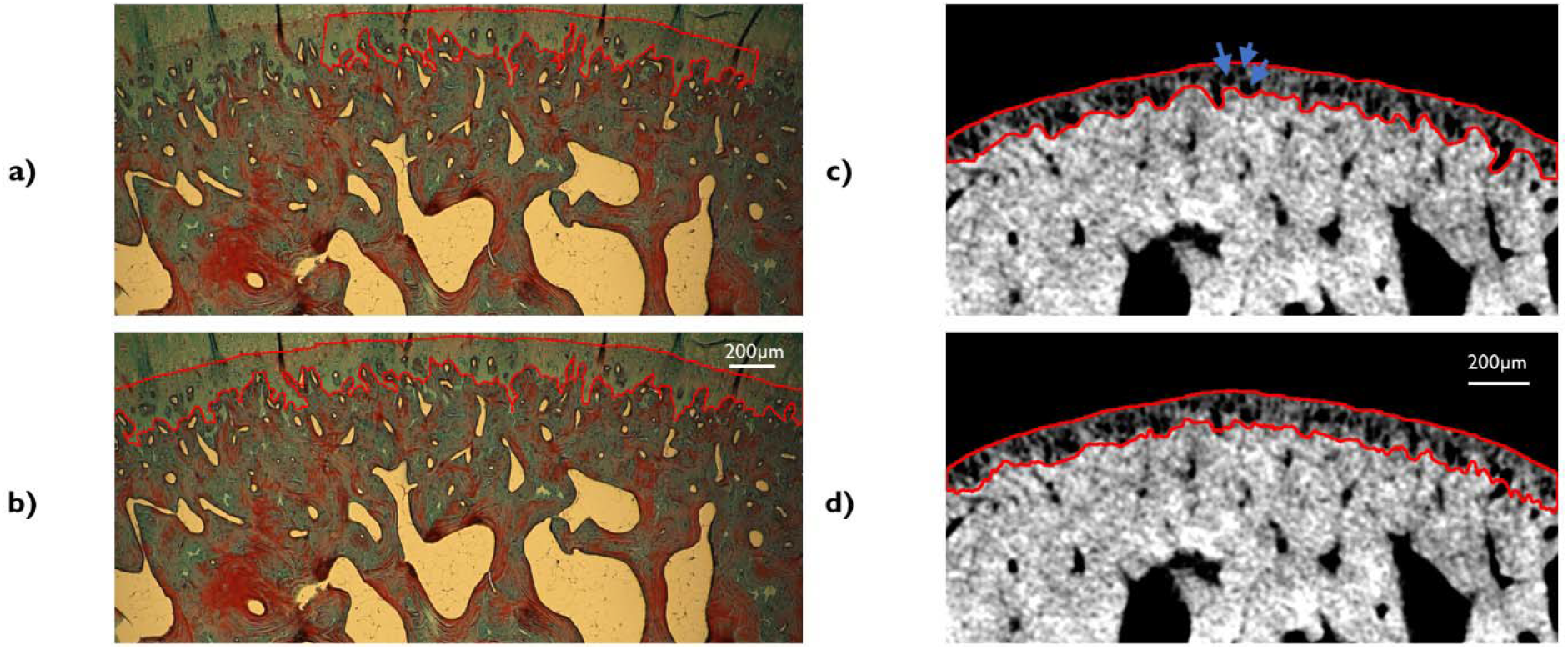
A histological section from the rabbit femoral condyle segmented manually (a) and automatically with the neural network (b). μCT image from a similar anatomical region (different animal) with manual (c) and automatic (d) segmentation. Scale bar for 200μm is shown in the corresponding images. The blue arrows refer to chondrocytes inside the CC layer.

The fully automatic CC segmentation was conducted using a deep learning pipeline inspired by *Solovyev et al.*^(40)^ on Python 3.7. The pipeline was built using in-house developed *Collagen*-framework (https://github.com/MIPT-Oulu/Collagen). For the histology segmentation, we used ResNet-34^(38)^ pre-trained on ImageNet^(34)^. We used a U-Net decoder with batch normalization in this model. The network was trained for 100 epochs under 4-fold cross-validation, splitting the training and validation folds with respect to rabbit ID. For the μCT segmentation, we used ResNet-18 as our base model, and also an FPN decoder, which had instance normalization as well as the spatial dropout. Briefly, the normalization reduces bias for individual features with large values, while dropout reduces model overfitting by zeroing random nodes of the network. This model was also trained in a 4-fold cross-validation but for 60 epochs due to faster convergence.

We used a combination of binary cross-entropy and soft Jaccard index as the optimization loss function. Binary cross-entropy is one of the most popular segmentation metrics and can result in stable convergence. However, Jaccard index can account for class imbalance, such as imbalance between the CC and the surrounding tissue. To facilitate a robust segmentation model, we used several image augmentation techniques (Supplementary Table 1) from the SOLT library^(42)^ to diversify the training data. To assess the final segmentation performance, we calculated the loss and Dice score coefficient as an average from the evaluation folds. The selection of the encoder and decoder was done based on an ablation study (Figure 2, Supplementary Figure 2).

**Figure 2.**
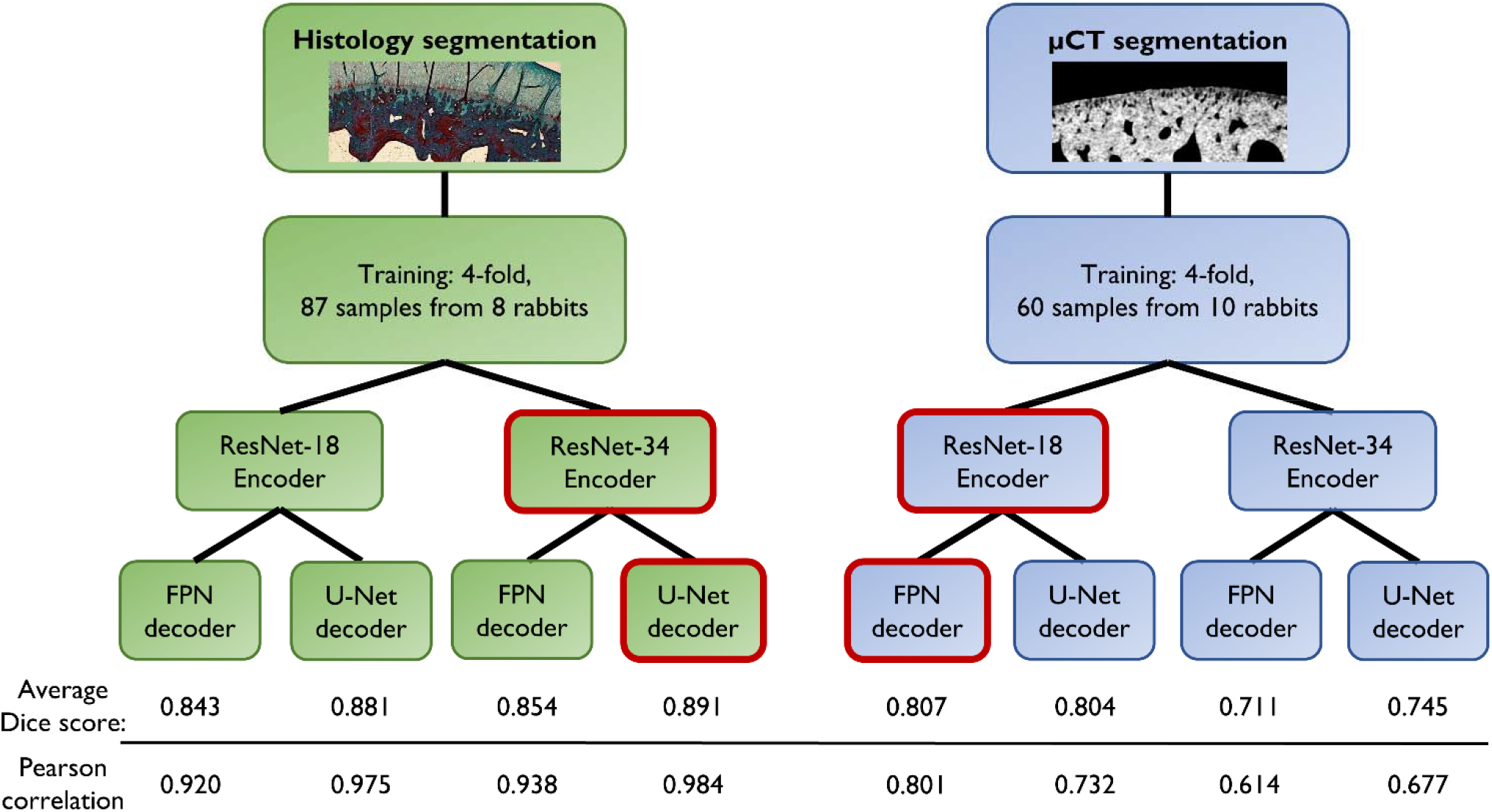
Illustration of the model training process. For both histology- and μCT segmentation, a total of four models were trained with two different encoder and decoder designs. Based on the experiments, ResNet-34 and U-Net were more suitable for the complex histology masks (Dice score = 0.891), while ResNet-18 and FPN yielded higher performance for the smoother μCT masks (Dice score = 0.807). Pearson correlation of the subsequent CC.Th analysis (bottom row) supported the choice of the segmentation models.

### Model application on new images (inference)

During inference, CC was predicted for the full histology images, by combining smaller tiles with a sliding window (512 × 1024 -pixel window with 256 × 512 -pixel steps), averaging the overlapping predictions. The tiling was used to avoid memory issues on the graphical processing unit while segmenting larger areas of CC. The tiles were combined, averaging the overlapping areas and predictions from every fold. Subsequently, a threshold was applied to the prediction map by using a probability of 0.8 (a high threshold was used for the exclusion of ambiguous areas from the maps, especially for the μCT images). In the case of the μCT stacks, the inference was conducted slice-by-slice with similar tiling. The predictions were averaged from every fold as well as the coronal and sagittal planes for obtaining the final probability map.

The histology masks were post-processed by removing small isolated areas (< 500 pixels). This ensured the removal of small artifacts while retaining large CC regions that could be disconnected due to a fold in the histology section (Supplementary Figure 3). In the μCT post-processing, masks were subjected to a sweep operation to keep only the largest object. This ensured the removal of possible false positives occurring on the tiles far from the actual CC layer. Finally, all CC masks were median filtered with a radius of 12 pixels (3D filtering in case of μCT).

### Morphological analysis

The full analysis procedure of CC.Th is summarized in Figure 3. The thickness estimation of the CC layer was performed automatically using a Python-based implementation of the local thickness algorithm. In the 2D case, the thickness assessment relies on mask skeletonization, a Euclidean distance transformation, and finally a simple circle-fitting algorithm^(43)^. The 3D CC.Th analysis of the μCT volumes was conducted with a similar sphere-fitting algorithm. From the estimated thickness maps, quantitative parameters such as mean-, median-, maximum CC.Th or standard deviation of CC.Th can be calculated. In this study, we used the mean CC.Th as the quantitative parameter. The source code for the full segmentation and analysis procedure is published on our research unit’s GitHub page (https://github.com/MIPT-Oulu/RabbitCCS). For the μCT volumes, the thickness analysis took 2-3 hours per sample (on a high-end 12-core CPU), whereas the analysis for the histology slices took roughly three seconds per image. For this study, the 3D thickness analysis was calculated with parallel processing on the Puhti supercomputer (https://research.csc.fi/csc-s-servers). This effectively reduced the computation time for the μCT volumes to roughly six minutes per sample.

**Figure 3.**
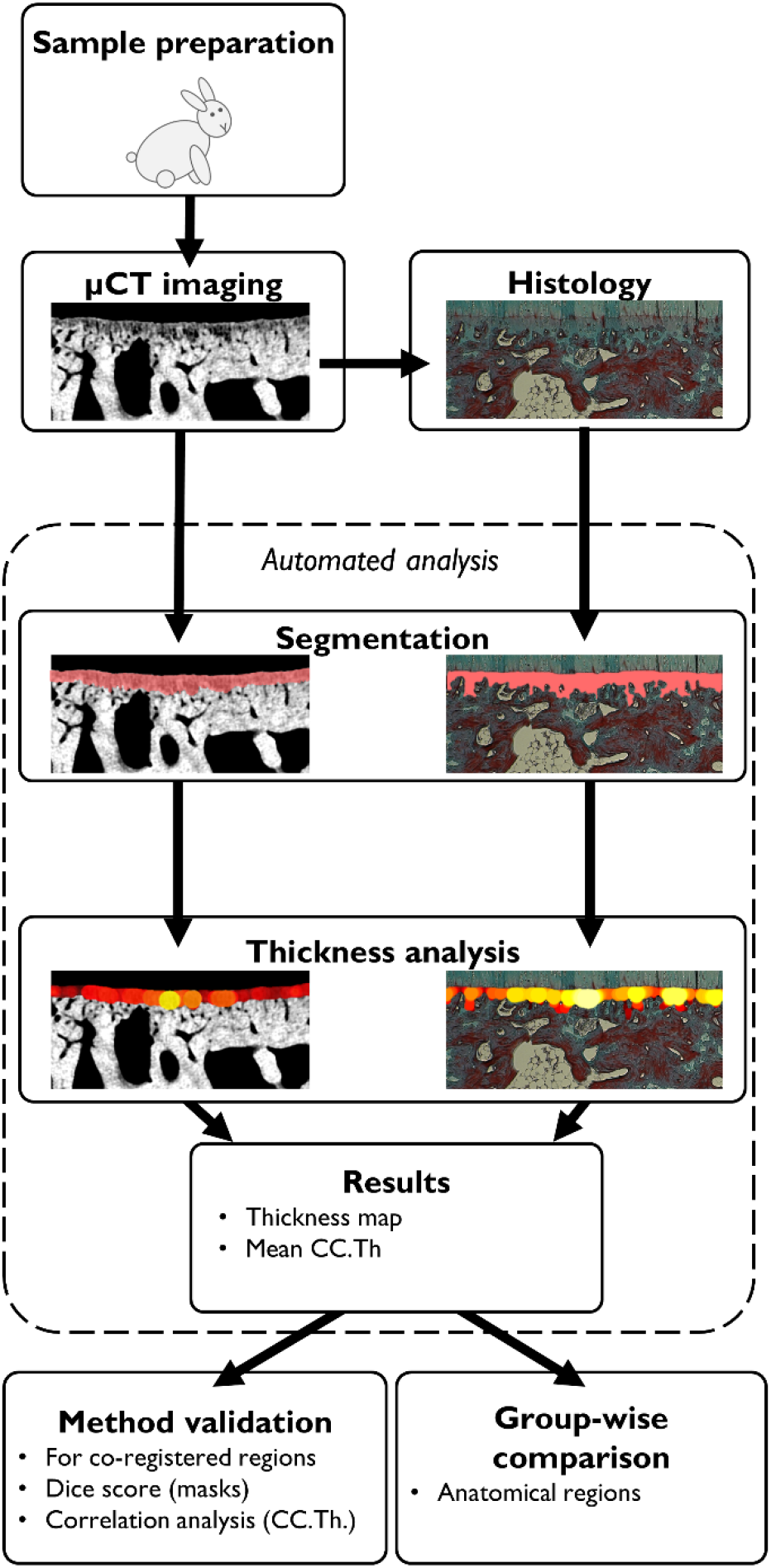
A flowchart summarizing the present study. After sample preparation, the tissue samples were imaged with μCT. Subsequently, the samples underwent histology processing, sectioning, and imaging with a light microscope. The preprocessing steps for the μCT data are illustrated in Supplementary Figure 1. During the automated analysis process, the CC layer is predicted using the deep learning models, thickness analysis is conducted, and finally, quantitative parameters are estimated from the estimated thickness maps. The obtained values were used in the validation of the methods as well as for comparison between the anatomical regions of the knee.

To further investigate the applicability of the automatic segmentation on CC.Th analysis, a 2D analysis was performed between the manual segmentations and the out-of-fold predictions of the selected models. The thickness values were averaged for each sample with multiple histology sections or μCT slices.

### Validation with histology

To compare the CC analysis between histology and μCT in 2D, matched μCT slices (Figure 4) were estimated using co-registration based on rigid transformations with DataViewer (Bruker, Kontich, Belgium; version 1.5.2.4). A total of 24 samples (from four animals) were co-registered with the corresponding histology sections to find the matching subchondral structures. Since the search space is large when aligning the few μm thick histology sections with the full sample, the remaining samples in paraffin blocks were imaged again using the μCT scanner. The co-registration of two μCT-imaged samples is straightforward and allows for locating the cutting orientation and approximating the location of the histological sample. Final co-registration was fine-tuned by performing a second co-registration between the original μCT datasets and the histology images. Five serial μCT images closest to the co-registered histology image were selected. Finally, we calculated the CC.Th from the co-registered histology image, while the CC.Th for μCT-imaged samples was averaged from the five selected images.

**Figure 4.**
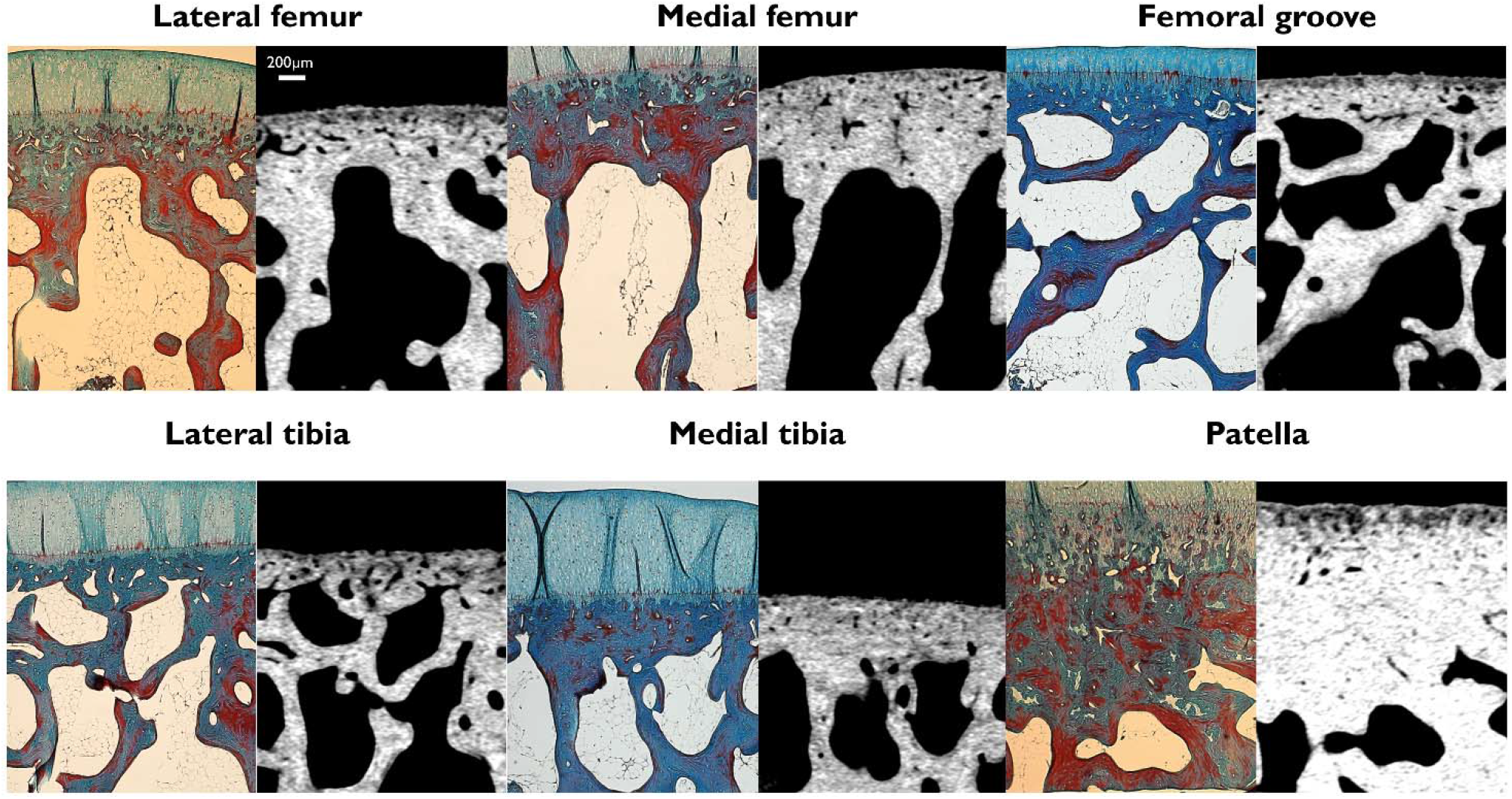
Examples from the co-registered histology slices and μCT images. Scalebar for 200μm is shown in the top left. The CC can be assessed using both imaging modalities, although the thinnest CC areas are not visible in the μCT images. Likely, these areas have a similar level of mineralization as the subchondral bone.

### Statistical analysis and performance evaluation

For the co-registration experiment, a two-tailed Pearson correlation and Bland-Altman analyses were conducted to compare CC.Th between the μCT and histology. The deep learning segmentation models were validated against the manual CC segmentations from μCT and histology using the Dice score. The thickness analyses using out-of-fold predictions and manual segmentations were compared using Pearson correlations. The anatomical differences of CC.Th were assessed using mean comparisons with Linear Mixed Effect Models, accounting for the rabbit ID as the random effect, and the anatomical location as the fixed effect. The significance was assessed with Least Significant Difference without Bonferroni correction.

## Results

### Deep learning-based segmentation

For both imaging modalities, the quality of the deep learning model predictions against the manual annotations (out-of-fold validation) is summarized in Figure 2 and Supplementary Figure 2. By comparing the four different model architectures, ResNet-34 with the U-Net decoder yielded the highest mean Dice score for histology (Dice score = 0.891), while ResNet-18 with FPN yielded the best performance for μCT segmentation (Dice score = 0.807). The quality of the segmentation on the full dataset was visually confirmed from virtual sections on orthogonal planes (Supplementary Figure 4).

In addition, we compared the 2D CC.Th analysis for the manual and predicted CC segmentations for both modalities (Figure 2 bottom, Supplementary Figure 5). With the selected model architecture, a high Pearson correlation was achieved between the manual and automatic CC.Th quantification from histology (*r* = 0.984). The correlation between predicted CC.Th and manually segmented CC.Th in μCT images was also strong, although considerably smaller (*r* = 0.801). This correlation analysis further supported the choice for model architecture (Figure 2, bottom)

### Validation with histology

Examples of μCT images co-registered with histology are shown in Figure 4. The results of the quantitative comparisons are shown in Figure 5 (predicted CC) and Supplementary Figure 6 (manual segmentation). The automated μCT-based measurements of CC.Th had a strong correlation (*r* = 0.897) with a similar analysis on the co-registered histology images. Furthermore, the μCT analysis had a good agreement (bias = 21.9 μm, standard deviation = 21.5 μm) with histology, based on the Bland-Altman analysis. Manual segmentation yielded a smaller correlation (*r* = 0.852) as well as greater bias (36.9 μm) and standard deviation (30.9 μm) than the comparison using predicted masks.

**Figure 5.**
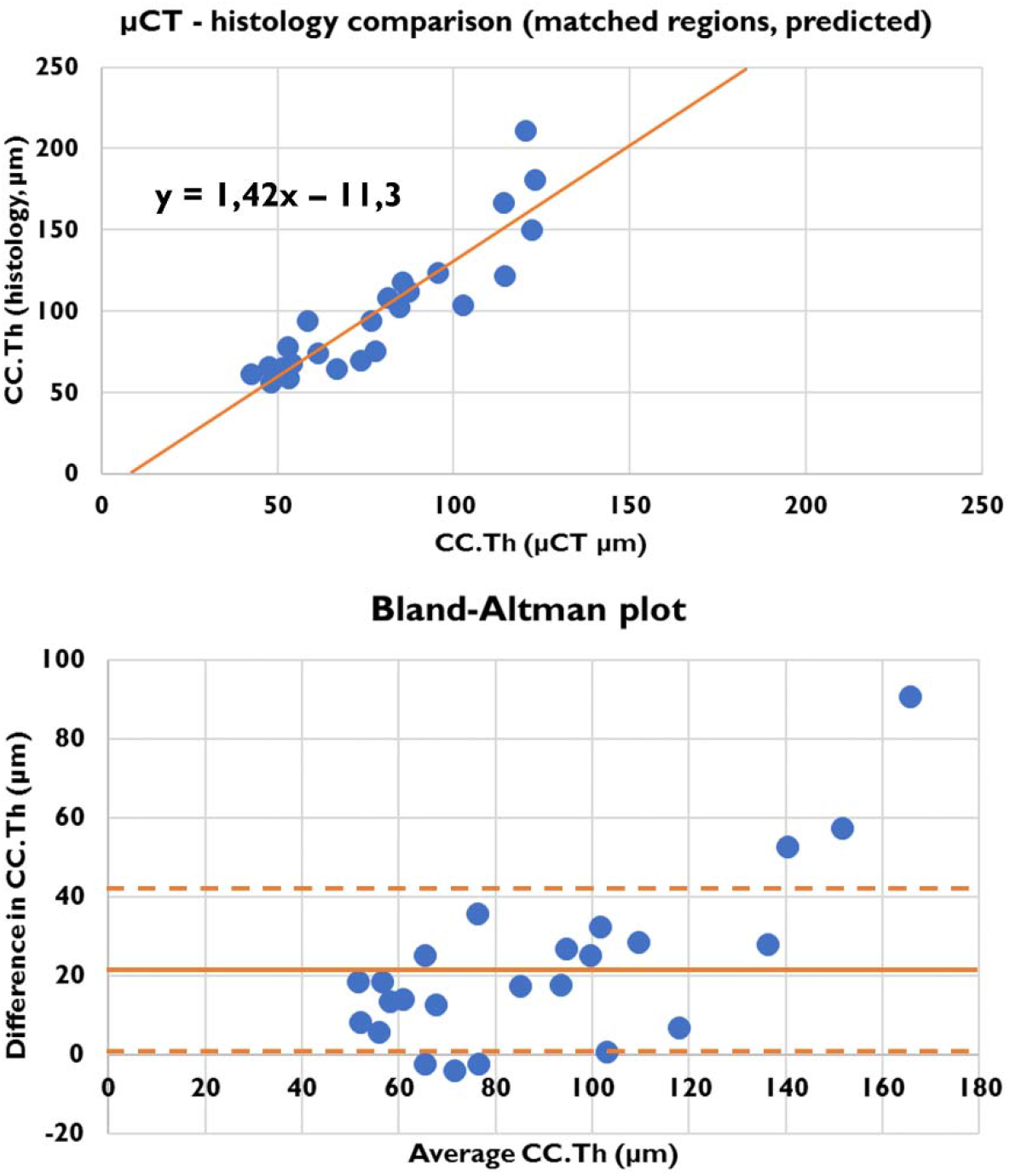
Quantitative CC.Th comparison of the matched histology and μCT regions based on automated segmentation. The equation for the linear fit is shown in the top image. For the Bland-Altman plot, the bias is indicated with a horizontal line, and the distance of one standard deviation with a dashed line. The estimated values are highly correlated (r = 0.897) and the Bland-Altman analysis reveals that the μCT method yields 21.9μm thinner CC.Th on average. The areas with a high CC.Th (mainly the patellar region) have the highest absolute differences between methods.

### Anatomical locations

An example of a thickness map and VOI inside a lateral plateau sample is shown in the Supplementary Video. The differences in CC.Th based on anatomical variability are illustrated in Figure 6. According to the Linear Mixed Effects Model analysis on the histology and μCT results (Table 2), the mean CC.Th varies greatly between the studied anatomical regions (*p* < 0.001). The thickest CC was in the patellar region, while the thinnest CC was in the tibial regions (lateral and medial plateau). The histology analysis allowed for further separation of the lateral and medial femoral condyles (*p* = 0.026). Although the absolute differences in CC.Th were larger using histology analysis than with the μCT approach, the μCT results had a smaller variance for individual regions than that observed with histology, allowing for separation of the anatomical locations.

**Figure 6.**
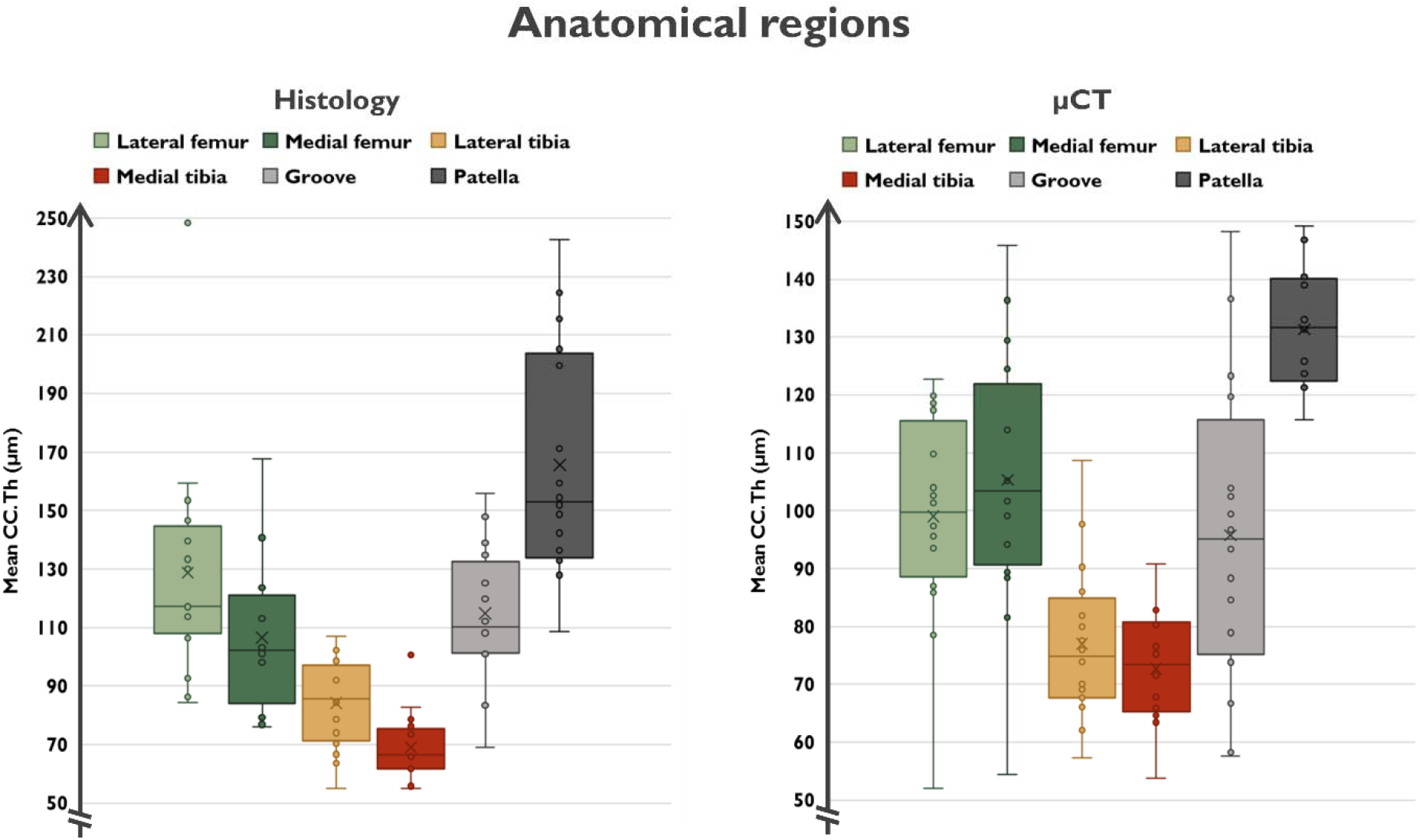
Boxplots illustrating the group-wise CC.Th values obtained from the histology and μCT modalities. The median value for each group is shown with the horizontal line and mean value with the cross. From the graph, the anatomical regions can be divided into three categories: thin CC (lateral and medial tibia), intermediate CC (lateral and medial femoral condyles, femoral groove) and thick CC (patella).

**Table 2.** Mean differences of mean CC thickness between the six regions (in μm): Lateral (LF) and medial (MF) femoral condyle, lateral (LP) and medial (MP) tibial plateau, femoral groove (G) and patella (P). The differences were assessed using a Linear Mixed Effects Model analysis, with Least Significant Difference. Detailed p-values are shown for p ≥ 0.001.

| **Histology** |  |  |  |  |  |
| --- | --- | --- | --- | --- | --- |
|  | MF | LP | MP | G | P |
| LF | **22.2**<br>**p = 0.026** | **44.5\*** | **59.7\*** | 13.9<br>p = 0.162 | **-36.9\*** |
| MF |  | **22.2**<br>**p = 0.026** | **37.4\*** | -8.4<br>p = 0.397 | **-59.1\*** |
| LP |  |  | 15.2<br>p = 0.126 | **-30.6**<br>**p = 0.002** | **-81.4\*** |
| MP |  |  |  | **-45.8\*** | **-96.6\*** |
| G |  |  |  |  | **-50.8\*** |
| **$\mu\text{CT}$** |  |  |  |  |  |
| LF | -6.3<br>p = 0.317 | **22.1**<br>**p = 0.001** | **26.3\*** | 3.3<br>p = 0.604 | **-32.3\*** |
| MF |  | **28.4\*** | **32.6\*** | 9.6<br>p = 0.131 | **-26.0\*** |
| LP |  |  | 4.2<br>p = 0.503 | **-18.8**<br>**p = 0.004** | **-54.4\*** |
| MP |  |  |  | **-23.0\*** | **-58.6\*** |
| G |  |  |  |  | **-35.6\*** |

\* $p < 0.001$

## Discussion

Morphological analysis of CC may reveal novel understanding of musculoskeletal physiology and pathology. A suitable tool for structural analysis of CC would be μCT, however, the separation between bone and CC is extremely challenging. In this study, we developed a μCT-based framework for 3D analysis of CC morphology. The framework utilizes state-of-the-art deep learning segmentation and automated analyses of CC.Th. Finally, we compared CC morphology on different joint surfaces within the healthy rabbit knees. Our results demonstrate that CC.Th can be quantified not only from histology but also from μCT, which is feasible and efficient due to an automatic segmentation approach. The proposed method enables studying the 3D morphology of the mineralized CC without the time-consuming and destructive histological processing and with minimal user-induced bias.

Our results revealed that different CNN architectures were best suited for CC segmentation from histology and μCT images. The FPN decoder is computationally more efficient, but it introduces an up-sampling layer for the model output. As a result, U-Net provides more detailed predictions since the CC is predicted without a subsequent interpolation. The results show that the U-Net decoder provided a slight advantage for segmenting the more complex CC structures in histology images. In the μCT images, such details are not visible, and FPN decoder yielded better results than the U-Net one. Encoder-wise, the deeper ResNet-34 might yield even better performance than the ResNet-18 encoder^(38)^. However, the ResNet-18 encoder with fewer layers than ResNet-34 performed better on the μCT data than ResNet-34. Thus, we suspect that the more complex ResNet-34 may overfit when images become ambiguous, as in the case of the μCT images.

The automated CC segmentation performed particularly well for the histology samples. A relatively high Dice score coefficient (0.891) and similar CC.Th results compared to the manual annotations (*r* = 0.984) suggest that the automated and manual methods give virtually identical results. For the μCT data, the performance was weaker than for the histology data (Dice = 0.807, *r* = 0.801). However, the segmentation of CC from the μCT images is much more difficult than segmentation from histology slides. Therefore, this result was expected. Based on our experience, there is also a significant variation in manual CC segmentation between human annotators. However, when comparing the estimated 2D CC.Th between histology and μCT for co-registered regions, there was strong agreement (*r* = 0.897).

We have previously shown that the subchondral bone plate imaged with μCT contains also the CC layer^(29)^. Consequently, automated labeling of the CC layer could identify the true subchondral bone tissue accurately. The proposed method requires high-resolution for resolving the mineralized cartilage. We believe that this is of high interest for studies that focus on the subtle changes in the bone plate, such as thinning due to increased remodeling. Such thinning of the bone plate has been suggested to occur already in the early stages of OA^(44)^.

The greatest differences in CC.Th between imaging modalities were seen in the samples with the highest CC.Th (Figure 5). We hypothesize that the less mineralized CC measured with μCT accounts for “young” tissue, which has distinct attenuation properties compared to the bone layer. Lower mineralization (hydroxyapatite content) of CC compared to bone has previously been reported using X-ray diffraction^(45)^. However, many studies have reported a higher mineralization of CC in backscattered electron imaging studies^(23,24,46)^, at least for human tissue. Thus, we considered that there might be a possible contribution of partial volume effects related to cellularity. Most of the cellular structures are visible, however, the observed changes are likely related to tissue mineralization. On the other hand, the deep CC appears more mineralized with similar attenuation properties as the subchondral bone, making it impossible to identify it solely based on X-ray methods. The samples with high CC.Th likely contain large, partly ossified areas of deep CC (such as the patellar region in Figure 4), leading to differences in CC.Th between the imaging methods. Therefore, we propose that our method could provide novel 3D information on tidemark advancement and other dynamic processes in calcified cartilage.

Interestingly, CC.Th depends greatly on anatomical location, as identified with both imaging methods. This is also consistent with our hypothesis. In the patellar region, CC.Th was the thickest among all locations of the rabbit knee. Femoral regions had intermediate CC.Th, while the thinnest regions were found in the medial tibial plateau region. We hypothesize that these variations in CC.Th are due to the distinct biomechanical environment in the different regions. First, the tibial plateau predominantly experiences compressive load due to body weight, while the patella experiences mainly shear forces that arise from the sliding joint articulation. Second, in the femur, the environment is a mix of these phenomena, i.e. the femoral condyles experience more compressive stress compared to higher shear forces on the groove. However, we did not find statistically significant differences in CC.Th between the condyles and groove. Finally, the higher shear stress experienced by the patella and femoral groove likely requires a stronger connection between the articular cartilage and the underlying subchondral bone plate, thus, resulting in higher CC.Th. Other studies have shown that the CC.Th of rabbit knees increases when subjected to chronic compression and that the CC is thicker in the lateral compared to the medial knee compartment^(47)^.

This study has several limitations: First, the decalcification process required for preparation of the histology slides may cause structural alterations in the tissue. Second, the intensity gradient between CC and subchondral bone can be ambiguous. This is especially the case for ultra-thin or non-existent CC. An ambiguous interface may appear because of endochondral remodeling resulting in bony protrusions into CC. Third, although an acceptable performance was achieved, the amount of training data used for the deep learning segmentation was relatively low. Examples from a greater number of animals may give a better performance, especially in the case of the challenging μCT segmentation. Fourth, our automated thickness analysis method is computationally expensive and does not scale well for large volumes. For routine use, more advanced scalable algorithms should be implemented, for example utilizing a distance ridge calculation^(48)^. Finally, the segmentation models might require fine-tuning to data acquired from a different microscope or μCT scanner to ensure sufficient performance on new samples.

In conclusion, we have presented a promising method for the morphological analysis of CC with μCT. To the best of our knowledge, this is the first automated method for quantitative 3D analysis of CC.Th that has been sufficiently validated against the histological gold standard. As a proof of concept, we could detect anatomical variation in the rabbit knee; the patellar region has the thickest CC and the tibial plateau region the thinnest. This structural difference between regions is presumably related to the diverse biomechanical environments, and thus the different requirements of the joint surfaces in different regions of the knee.

## Acknowledgements

The full source code of the project is openly available on our research unit’s GitHub page (https://github.com/MIPT-Oulu/RabbitCCS). This project received funding from the European Union’s Horizon 2020 research and innovation programme under the Marie Skłodowska-Curie grant agreement No 713645; European Research Council under the European Union’s Seventh Framework Programme (FP/2007-2013)/ERC Grant Agreement No. 336267; Academy of Finland [grant numbers 286526, 303786, 324529]; Saastamoinen Foundation, Päivikki ja Sakari Sohlberg Foundation; Finnish Cultural Foundation (Central Fund No. 191044, North Ostrobothnia Regional Fund No. 60172246); Maire Lisko Foundation; The Canadian Institutes of Health Research, The Canada Research Chair Programme; The Killam Foundation. The strategic funding of the University of Oulu and the University of Eastern Finland are acknowledged. CSC – IT Center for Science, Espoo, Finland is acknowledged for generous computational resources.

## Author’s roles

Study conception and design: SJOR, LH, PT, AT, RKK, SS, WH, MAJF

Data collection: LH, PT, RKK, WH, MAJF

Method development: SJOR, AT, EP, SS, MAJF

Data analysis and interpretation: SJOR, LH, AT, SS, MAJF

Drafting the manuscript: SJOR, LH, MAJF

Critical revision and approving the final version of the manuscript: All authors

SJOR takes responsibility for the integrity of the work.

## Supplementary material

**Supplementary Figure 1.**
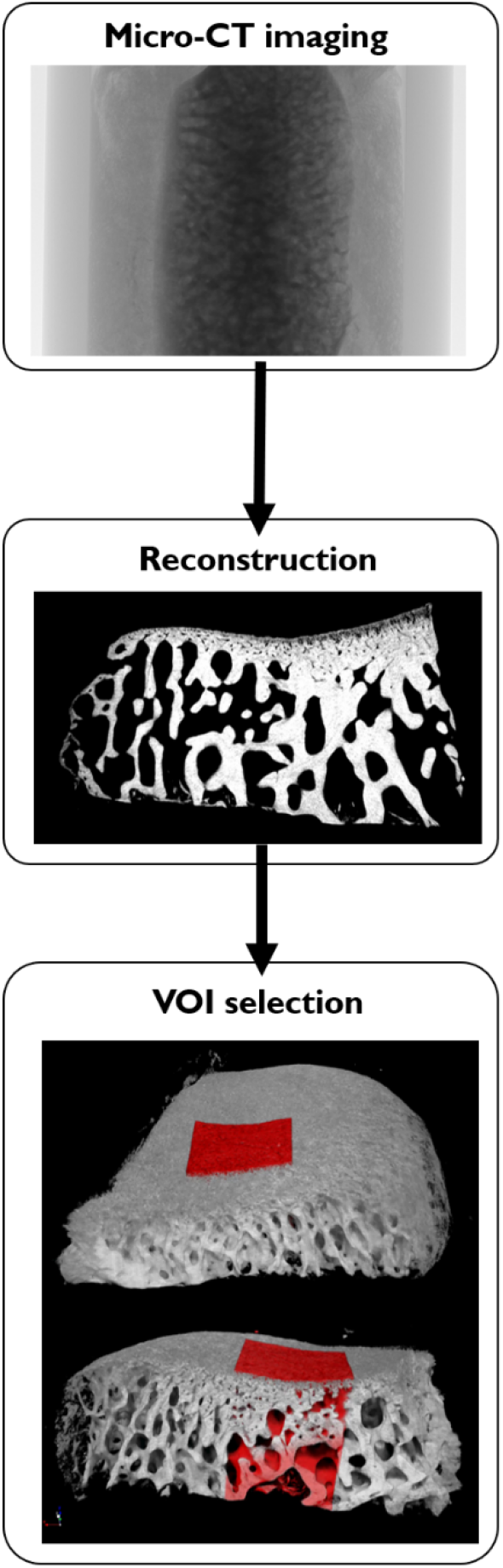
The main preprocessing steps for the μCT data. In the top image, an example projection image is shown from a lateral tibial plateau sample. In the middle, a coronal section from the reconstruction result is displayed. The bottom image shows the 3D rendered sample with an example of the VOI selection. The top part shows an overview, while the bottom part includes a virtual section inside the sample.

**Supplementary Figure 2.**
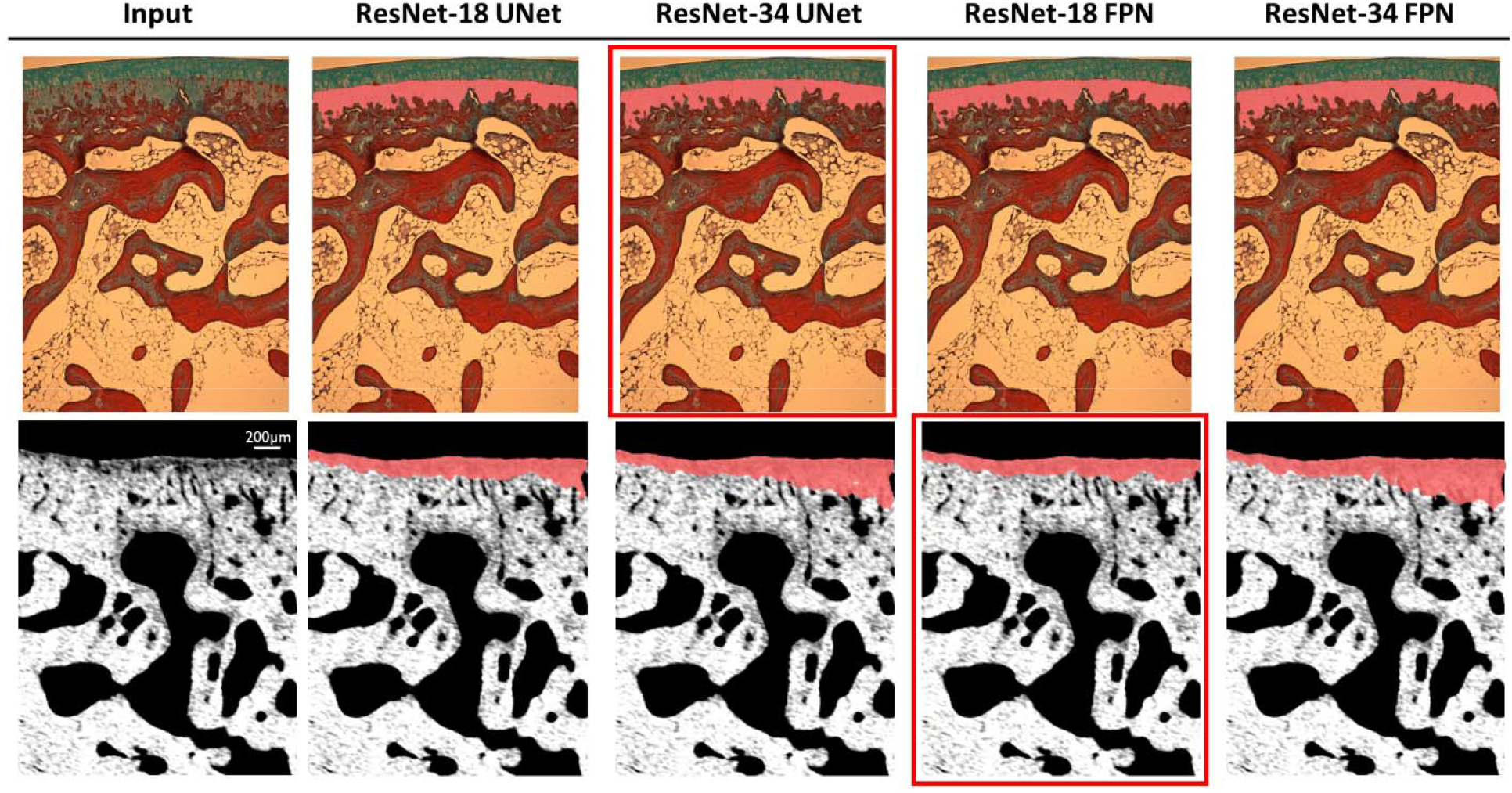
Qualitative comparison of the predicted CC masks. The selected models are highlighted with a red rectangle. Since the FPN decoder requires upsampling of the segmentation result, small details of the histology masks are easily overlooked. This makes it better suited for the smooth μCT masks. U-Net preserves the complexity of the histology mask slightly better than FPN. For the μCT models, the complex ResNet-34 model may overfit the training data, thus yielding higher errors on the validation images.

**Supplementary Figure 3.**
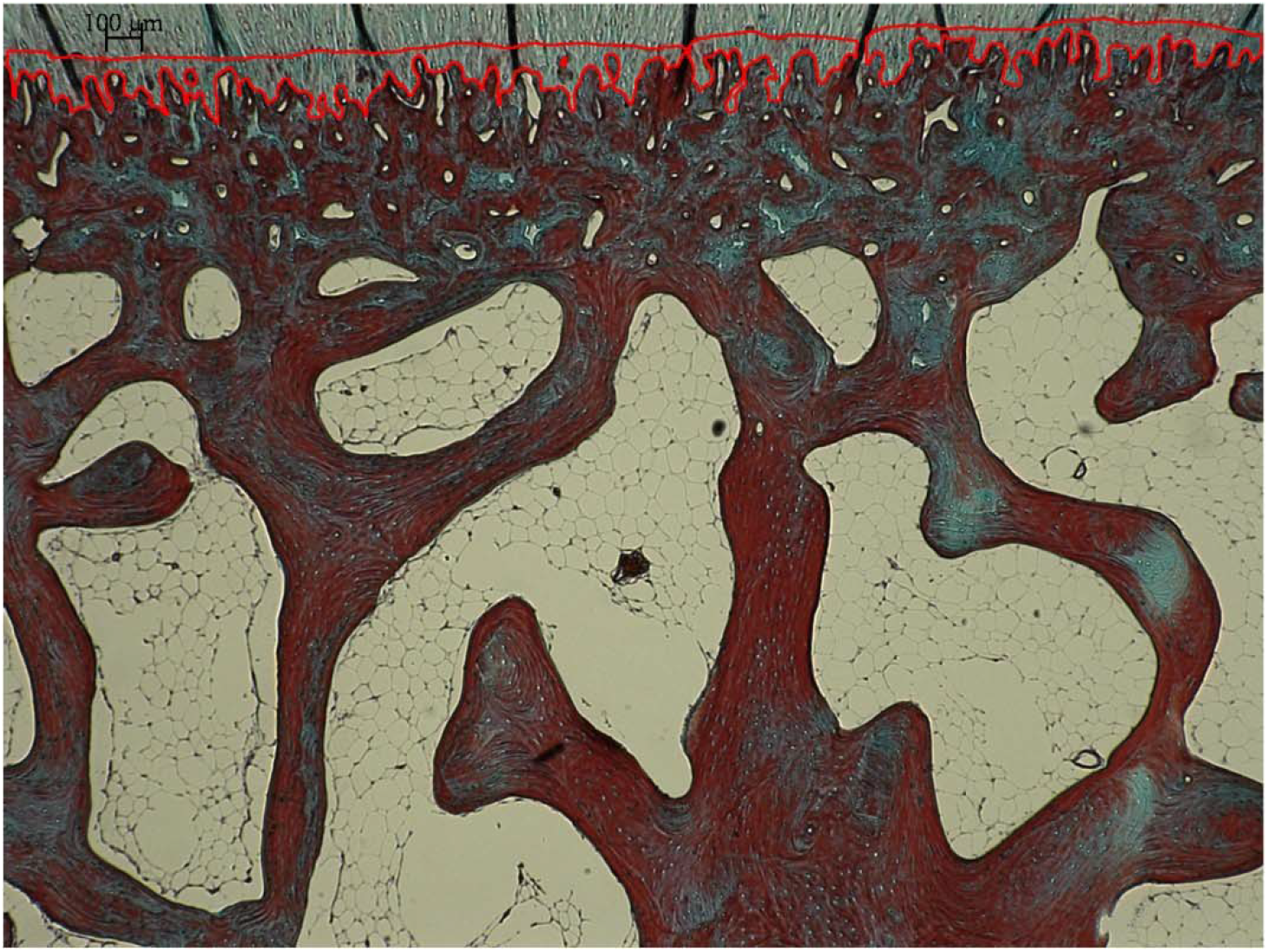
Example histology slice with a disconnected prediction. The red outline illustrates the mask post-processed with the despeckle operation, removing areas smaller than 500 pixels. If only the largest mask would be retained, much of the real CC layer would be lost.

**Supplementary Figure 4.**
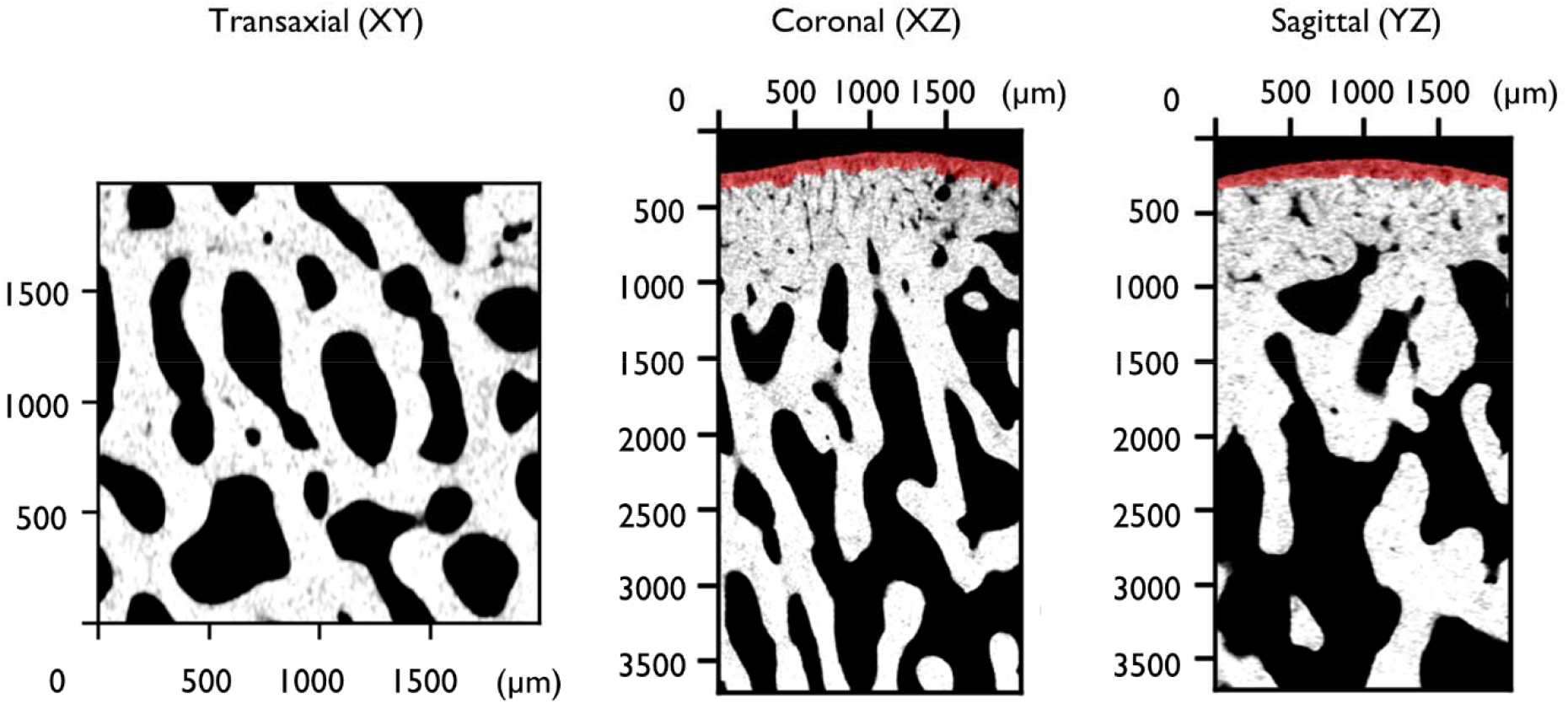
Illustration for visual output of the predicted CC mask. A figure with three orthogonal planes on the sample is drawn after the inference, displaying the model output in red.

**Supplementary Figure 5.**
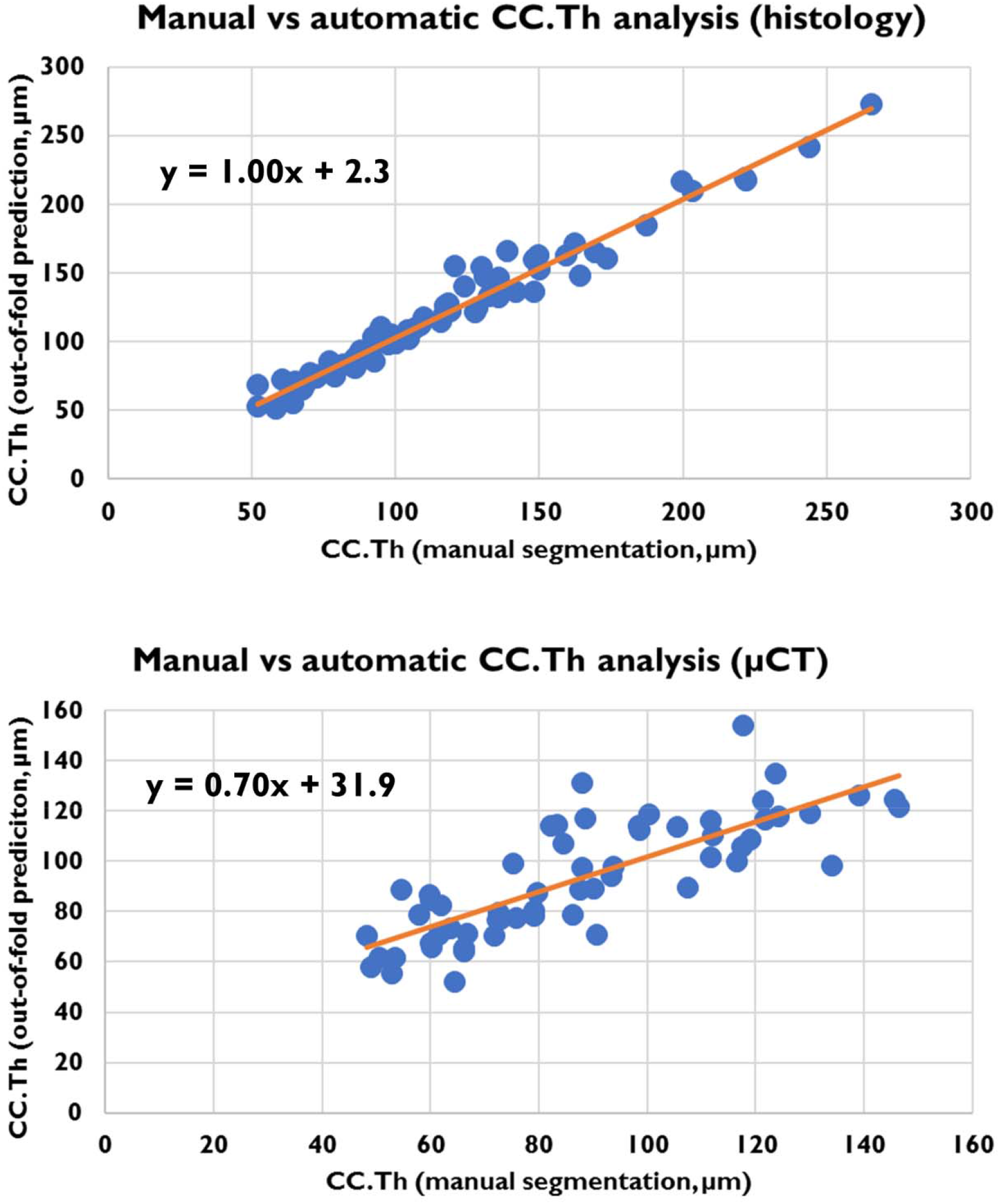
Scatterplots for the CC.Th analysis based on manual and automatic segmentation. The equation for the linear fit is shown. With the histology images, the correspondence between the methods is extremely high. The more challenging μCT segmentation results in discrepancies between prediction and the gold standard.

**Supplementary Figure 6.**
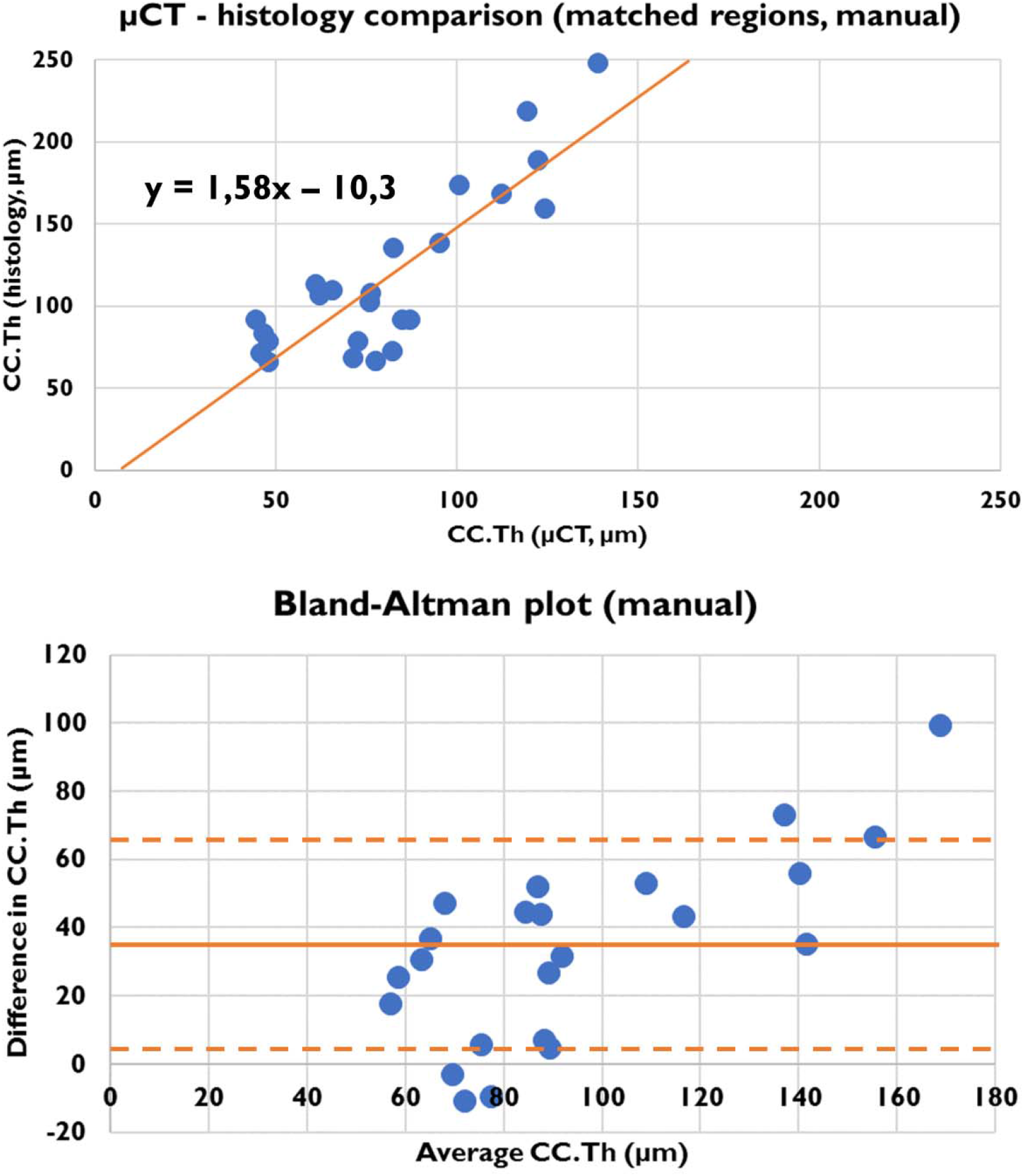
Quantitative CC.Th comparison of the matched histology and μCT regions based on manual segmentation (compared to the predicted CC in Figure 5). The equation for the linear fit is shown in the top image. For the Bland-Altman plot, the bias is indicated with a horizontal line, and the distance of one standard deviation with a dashed line. The analysis yielded a Pearson correlation coefficient of 0.852, a bias of 36.9μm, and a standard deviation of 30.9μm.

